# The *Caenorhabditis elegans* centrosome is surrounded by a membrane reticulum, the centriculum, that affects centrosome size and function

**DOI:** 10.1101/2022.06.15.496339

**Authors:** Richa Maheshwari, Mohammad M. Rahman, Seth Drey, Megan Onyundo, Gunar Fabig, Michael A. Q. Martinez, David Q. Matus, Thomas Müller-Reichert, Orna Cohen-Fix

## Abstract

Centrosomes are membraneless organelles that nucleate microtubules. At their core is a pair of centrioles that recruit pericentriolar material (PCM), a phase-separated condensate. In many cell types, including human cells, centrosomes are surrounded by endoplasmic reticulum-derived membranes of unknown structure and function. Using volume electron microscopy, we show that the *C. elegans* centrosome is surrounded by a membrane reticulum that we call the centriculum, for ***centr***osome-associated membrane ret***iculum***. Increasing centriculum size by genetic means led to expansion of the PCM and increased microtubule nucleation capacity, an unexpected finding given that the PCM is a membraneless condensate. We provide evidence that the centriculum serves as a microtubule “filter” by limiting the number of microtubules that can elongate fully. We also show the centriculum fuses with the nuclear envelope during mitosis. We propose that this fusion contributes to nuclear envelope breakdown by transducing forces from the elongating spindle to the nuclear membranes.

## Introduction

Centrosomes are membraneless organelles that serve as microtubule-organizing centers, such as at the poles of mitotic spindles. They are composed of a pair of centrioles surrounded by pericentriolar material (PCM) that promotes microtubule nucleation. The PCM increases in size as cells progress through the cell cycle, in a process known as centrosome maturation (reviewed in (Gomes Pereira et al., 2021)). During this process, recruitment of PCM proteins, such as the *C. elegans* SPD-5, the functional homolog of CDK5RAP2/Cnn, depends on the centrosomal proteins SPD-2/CEP192 and the Aurora and Polo-like kinases (Kemp et al., 2004; Pelletier et al., 2004; Pintard and Bowerman, 2019). The PCM, in turn, recruits proteins that promote microtubule nucleation, such as the γ-tubulin ring complex (γ-TuRC), and increases the local concentration of tubulin (Woodruff et al., 2017; Baumgart et al., 2019). The PCM in *C. elegans*, as well as human cells, has been suggested to form a liquid-liquid phase-separated (LLPS) condensate (Ahn et al., 2020; Jiang et al., 2021; Woodruff et al., 2017; Zwicker et al., 2014). As such, the size of the PCM is expected to be a function of the concentration of its components and the properties of the surrounding liquid environment (Shin and Brangwynne, 2017; Woodruff et al., 2017).

In most organisms, centrioles are provided by the sperm. Soon after fertilization, centrioles recruit the PCM and duplicate to form two centrosome, which remain associated with the male pronucleus via the linker of nucleoskeleton and cytoskeleton (LINC) complex (Malone et al., 2003; Meyerzon et al., 2009). The centrosomes move away from each other and will ultimately form the first mitotic spindle of the zygote. Centrosome separation is facilitated by interactions between microtubules nucleated by the centrosomes and the nuclear envelope (NE) (Blangy et al., 1995) in a process that is not fully understood. The NE itself is made of two membranes, the outer nuclear membrane and the inner nuclear membrane, that are traversed by nuclear pore complexes (NPCs) that allow transport of material between the cytoplasm and nucleoplasm (Cohen-Fix and Askjaer, 2017). After pronuclear meeting, as the embryo enters its first mitosis, nuclear membrane-associated proteins dissociate from the NE, and the nuclear membranes themselves become fenestrated in a process known as NE breakdown (Cohen-Fix and Askjaer, 2017). During this time, centrosomes start nucleating microtubules that extend towards the chromosomes and ultimately form the mitotic spindle.

In the *C. elegans* early embryo, centrosomes are surrounded by an ER-derived membrane ((Poteryaev et al., 2005) and Fig. 1A; ER is detected by the signal peptidase, SP12, fused to mCherry). Centrosome-associated membranes were also observed in other systems, such as *Drosophila*, sea urchin, PtK2 cells and LLC-PK2 cells (Bergman et al., 2015; Diaz et al., 2019; Harris, 1975; Karabasheva and Smyth, 2019; Waterman-Storer et al., 1993). The configuration of these membrane and their function are unknown. Using volume electron microscopy (vEM) analysis, we show that the membrane around the *C. elegans* centrosomes forms a reticulum, leading us to name it the “centriculum”, for **<u>centr</u>**osome-associated endoplasmic ret**<u>iculum</u>**. We provide evidence for an interdependent relationship between the PCM and the centriculum: the centriculum requires the PCM for its formation. Conversely, PCM size and its microtubule-nucleating capacity are affected by the size of the centriculum, an unexpected finding given that the PCM is a LLPS condensate. Our data also suggest that the centriculum acts as a “microtubule filter” by restricting microtubule elongation to those that are in the appropriate orientation to pass through the membrane reticulum. Finally, we propose that during mitosis, the centriculum plays a role in associating the centrosome with the NE, thereby promoting nuclear membrane fenestration.

**Figure 1:**
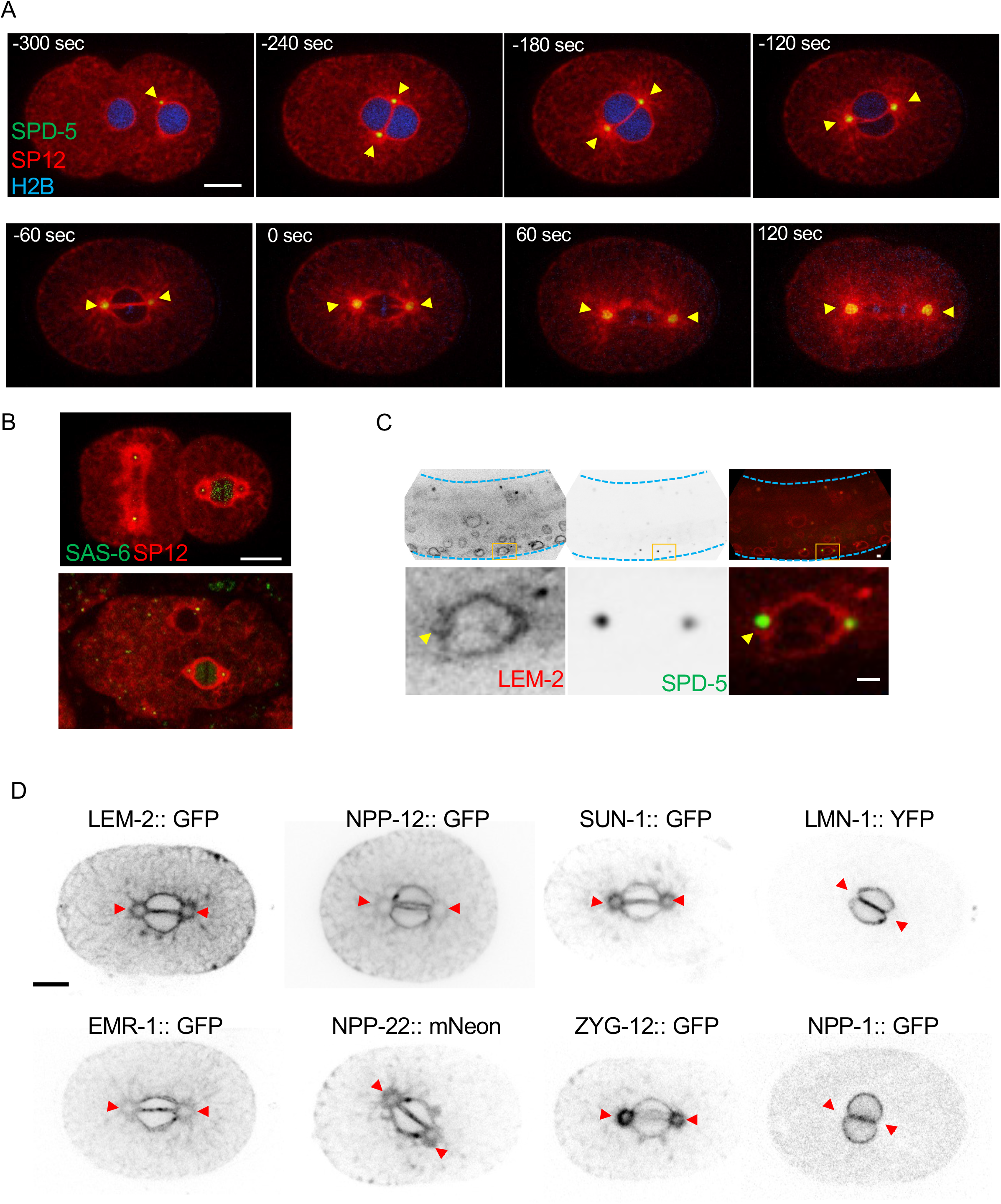
The centrosome is surrounded by ER membrane that contains some, but not all, NE proteins. (A) Time lapse images of a *C. elegans* one-cell embryo, expressing the resident ER signal peptidase, SP12, fused to mCherry (red), the PCM protein, SPD-5, fused to GFP (green), and histone H2B fused to CFP (blue) (strain OCF164). Yellow arrowheads point to the membrane surrounding the centrosomes. Time points are relative to metaphase (t=0), as determined by the appearance of the membrane gap at the membrane interphase between the two pronuclei (note that the histone H2B signal is faint and not always visible in all embryos). In the first time point (-300 sec), the pronucleus containing the sperm DNA and associated with the centrosomes is on the right. (B) A 2-cell embryo (top) and a multi-cell stage embryo (bottom) expressing the centriolar protein SAS-6 fused to GFP (green) and SP12 fused to mCherry (red) (strain OCF124). (C) A vulval precursor cell (VPC, bottom row) from a representative L3 stage *C. elegans* larvae (top row) expressing endogenously tagged NE/ER protein LEM-2 fused to mCherry (red) and PCM protein SPD-5 fused to GFP (green) (strain OCF173). The dashed blue lines in the upper panel outline the worm. The images on the bottom are an enlargement of the area marked in orange in the top panel. The yellow arrow points to the membrane around the centrosome (the second centrosome is on a different focal plane). (D) Localization pattern of the indicated NE proteins relative to the membrane around centrosomes (indicated by red arrowheads) in 1-cell embryos at prometaphase. Scale bar in all panels=10 μm

## Results

### The centrosome is surrounded by an ER membrane that contains some, but not all, NE proteins

After fertilization in *C. elegans*, chromosomes originating from the oocyte and sperm become encased in separate nuclear envelopes to form the two pronuclei. The pronuclei subsequently move towards each other and meet, forming a flat membrane surface at the interface between them. At the same time, the duplicated centrosomes increase in size by recruiting additional PCM material and separate via microtubule-associated forces until they reach opposite ends of the pronuclear membrane interface (Fig. 1A, -240 sec, reviewed in (Pintard and Bowerman, 2019)). Throughout this process, the centrosomes are surrounded by an ER-derived membrane system ((Poteryaev et al., 2005) and Fig. 1A), the configuration and function of which is unknown. Centrosome-associated ER membranes are also present in multi-cellular embryos (Fig. 1B) and in vulval precursor cells (VPC) (Fig. 1C). Thus, the association of centrosomes with ER membrane appears to be a general feature in *C. elegans*. Since the membranous structures surrounding the centrosome were the largest and most easily visualized in 1-cell embryos, we decided to further characterize them at this developmental stage.

The proximity of the centrosome to the NE led us to examine which NE-associated proteins also localize around the centrosome. We found that membrane-embedded NE proteins, such as the inner nuclear membrane proteins LEM-2 and EMR-1 (Emerin), the LINC complex proteins SUN-1 and ZYG-12, and the transmembrane NPC subunits such NPP-12 (human homolog: gp210) and NPP-22 (NDC-1), all localize around the centrosome (Fig. 1D). In contrast, the peripheral NE proteins LMN-1 and the NPC subunit NPP-1 (Nup54) did not localize around centrosomes (Fig. 1D). Published data indicate that other NPC subunits, such as NPP-3 (Nup205), NPP-19 (Nup35), and NPP-5 (Nup107), also do not localize around centrosomes (Hachet et al., 2012; Martino et al., 2017; Ródenas et al., 2009). Thus, membrane-embedded NE proteins localize to both the NE and around centrosomes, while peripheral NE proteins are confined to the NE. In the case of EMR-1, NPP-12 and LEM-2, the localization around the centrosome may have reflected a general ER localization. However, SUN-1 and ZYG-12 appear to be specifically enriched at centrosome-associated membranes during mitosis.

### The membrane around the centrosome forms a mesh-like, reticular structure, now named the centriculum

To determine the configuration of the membrane around centrosomes we used a volume electron microscopy (vEM) technique called Focus Ion Beam - Scanning Electron Microscopy (FIB-SEM, (Kizilyaprak et al., 2019; Narayan and Subramaniam, 2015; Rahman et al., 2020; Xu et al., 2017)). Briefly, individual 1-cell *C. elegans* embryos in mitosis were high-pressure frozen, followed by freeze substitution and embedding in resin. A trench was created near the area of interest, and an ion beam sequentially milled off 9 nm from the block, with a SEM image taken after each iteration (final voxel size= 9 nm^3^). In the present study, we present 3D reconstruction of FIB-SEM data from (Rahman et al., 2020) of membranes around three prophase centrosomes and three metaphase centrosomes.

In a single SEM slice from the vEM image stack, centrosomes appeared as an electron dense region at opposite poles of the interface between the two pronuclei (Fig. 2A, B, C, F, G, K and Supplemental Fig. S1 and S2, panels i and ii). In our staining method, which was optimized for maximal membrane contrast, microtubules were observed only occasionally (see (Rahman et al., 2020)) and centrioles were never detected. The need for three-dimensional information becomes apparent when examining a single EM image, where membranes around the centrosome could have been interpreted as an aggregation of vesicles (Fig. 2A). However, segmentation (i.e., 3D-reconstructions) of 400-500 nm above and below a center plane of centrosomes revealed that the membranes around centrosomes form a reticulum (Fig. 2C-E, G-I and Supplemental Fig. S1 and S2; segmentations were done to the edge of the reticulum, beyond which only ER tubules were present). This configuration was also confirmed by electron tomography data (see below). Based on this, we named the spherical membrane structure that surrounds centrosomes the centriculum, for **centr**osome-associated membrane ret**iculum** (plural= centricula). The configuration of membrane in the centriculum is distinct from the previously reported membrane configurations of the ER, typically sheets and tubules, or the NE, which is a membrane sheet (Zhang and Hu, 2016).

**Figure 2:**
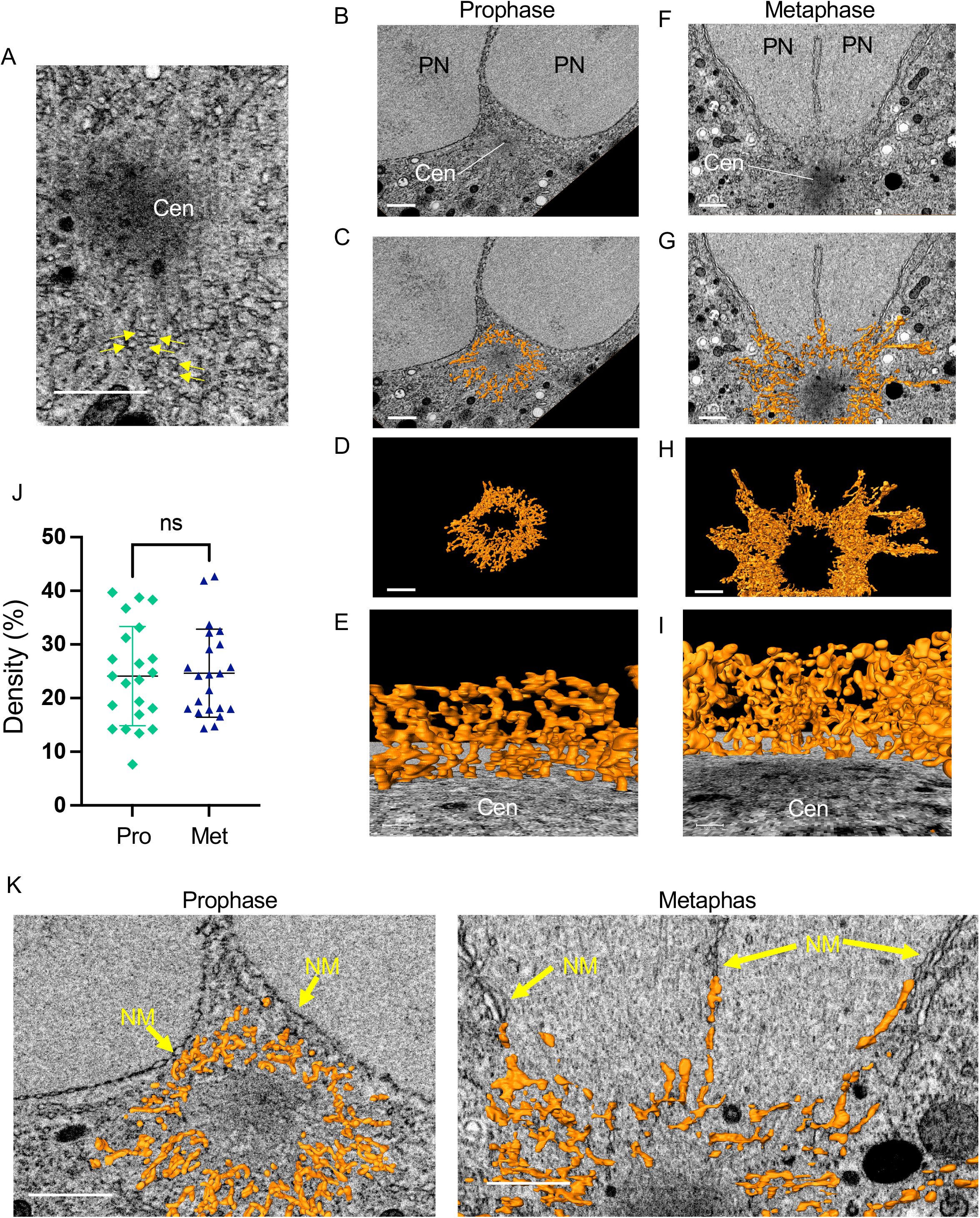
The membrane around the centrosome forms a reticulum. (A) A single SEM image from a region surrounding the centrosome in metaphase. Arrows point to membrane structures. The centrosome (Cen) appears as a dark zone that, when segmented using 3D FIB-SEM data, forms an irregular sphere (not shown). Scale bar= 1 μm. (B-I) 3D reconstruction using FIB-SEM data from (Rahman et al., 2020) of the membranes around the centrosome in a 1-cell embryo in prophase (panels B-E) and metaphase (panels F-I). Additional examples of prophase and metaphase centrosomes and their associated membranes are shown in Supplemental Fig S1 and S2, respectively. A portion of the two pronuclei (PN) is visible adjacent to the centrosome in panels B, C, F and G. (B, F) A SEM image from the centrosome’s mid-section. Scale bar= 1 μm. (C, G) The same SEM images as in panels B and F, superimposed with a 200 nm (in the z axis) of segmented membrane around the centrosome (orange). Scale bar= 1 μm. (D, H) An 800 nm segment of the reconstructed membranes surrounding the centrosome from the same prophase (D) and metaphase (H) embryos as above. Scale bar= 1 μm. (E, I) A view from the centrosome towards the pronuclei/chromosomes (not shown) through the entire centriculum wall. Scale bar = 100 nm. (J) Centriculum density was determine by cropping at least 10 consecutive 250 nm^3^ cubes from the center portion of two centricula in 1-cell embryos in either prophase (Pro) or metaphase (Met), and determining the percent volume that is occupied by a membrane. Bars indicate means and standard deviations. Statistical analysis was done using Student t-test. K. Enlarged images of the same centricula as in panels C and G, with a 100 nm slice of segmented membrane superimposed on the SEM image. Arrows point the nuclear membranes (prophase) or remnants thereof (metaphase).

The density of the centriculum was the same in prophase and metaphase, as determined by the amount of membrane per unit volume (Fig. 2J). However, the extent of centriculum association with the NE changed during the cell cycle. During prophase, the centriculum is distinct from the pronuclear membranes (Fig. 2K, left panel and Supplemental Fig. S1, Ai and Bi). In contrast, during metaphase the centriculum is indistinguishable from the remnants of the pronuclear membranes (Fig. 2K, right panel Supplemental Fig. S2, Ai and Bi), which remain as perforated sheets around and between both pronuclei (Rahman et al., 2020). This observation suggests that the centriculum may play a role in associating the centrosome with mitotic nuclear membrane remnants, a process that could implicate spindle elongation in nuclear envelope breakdown (see Discussion).

### Centriculum size is dependent on the integrity of the centrosome

Having established that the membrane around the centrosome, the centriculum, has a reticular structure, we next investigated the spatial and functional relationship between centrosome components and the centriculum. To this end, we first examined the localization of centrosomal proteins with respect to the centriculum at metaphase, starting with an inner most centrosomal protein (SAS-6, a centriole component) and working our way outwards (Magescas et al., 2019): the inner PCM protein, SPD-2, the outer PCM proteins, SPD-5, the Aurora A kinase, AIR-1, and finally alpha tubulin, TBA-2, to examine microtubules. SAS-6, SPD-2 and SPD-5 were clearly within the confines of the centriculum (Fig. 3A). The fluorescence associated with tagged AIR-1 and TBA-2 was less well demarcated (Fig. 3B), and we therefore quantified the fluorescence intensities of SP12 and AIR-1 or TBA-2 along a line that traversed the centriculum (Fig. 3C). Our data show that the centriculum is outside the peak intensities of both AIR-1 and TBA-2, suggesting that it surrounds the outermost part of the centrosome.

**Figure 3:**
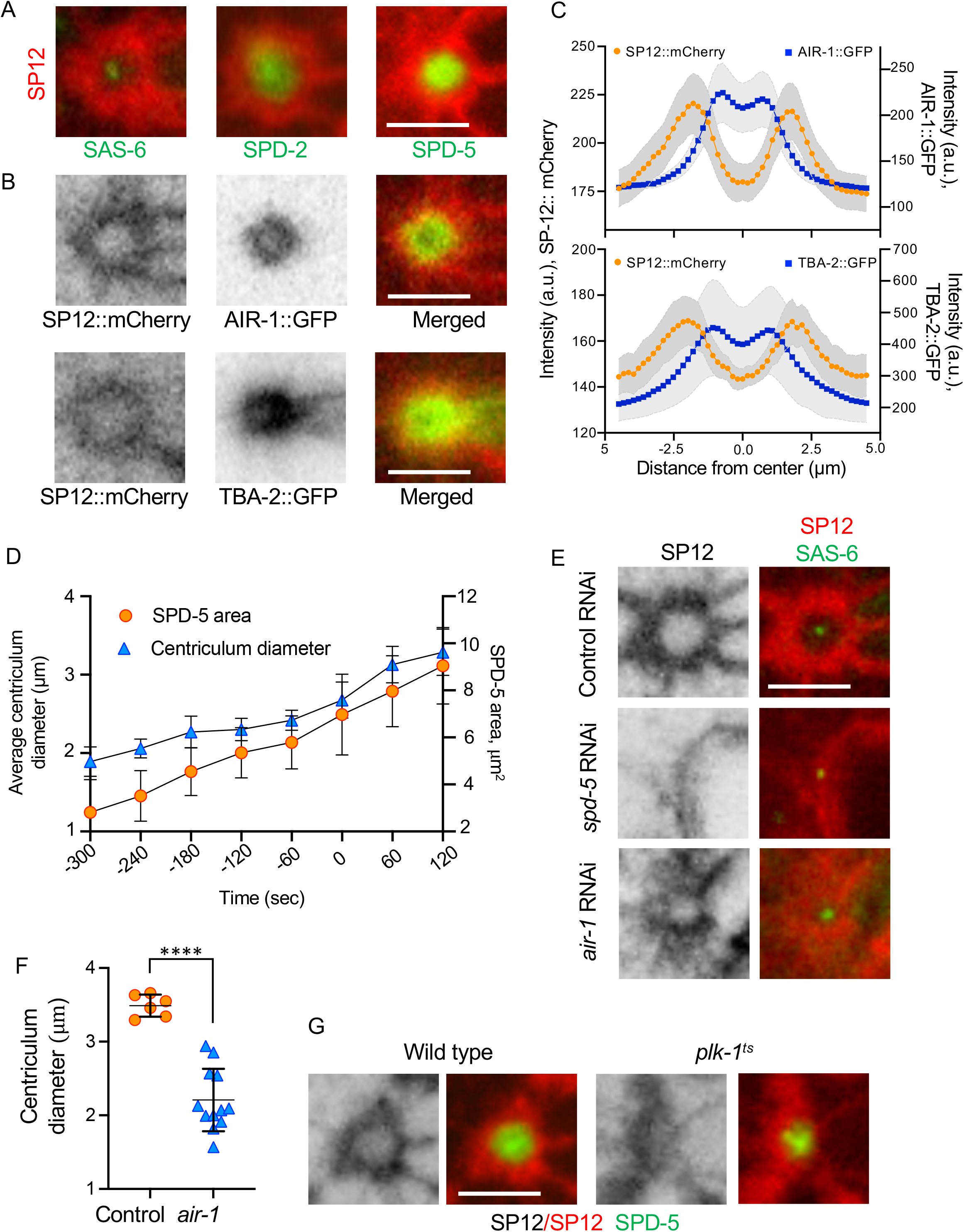
Centriculum size depends on the integrity of the centrosome. (A) The spatial relationship between the centriculum (detected using SP12::mCherry) and the following components of the centrosome in a 1-cell stage embryo at metaphase: SAS-6::GFP (strain OCF124), SPD-2::GFP (OCF127) and SPD-5::GFP (OCF164). Representative images are shown. Scale bar = 5 μm. (B) The spatial relationship between the centriculum (detected using SP12::mCherry) and either AIR-1::GFP (top row, OCF172) or TBA-2::GFP (bottom row, MSN146). Representative images are shown. Scale bar = 5 μm. (C) Fluorescence intensities along a line that traverses the centrosome and its surrounding centriculum using the same strains as in panel B (n=16 for the top panel and 8 for the bottom one). Error (in grey) around the mean (in orange for SP12::mCherry and blue for the GFP tagged protein) represents 95% confidence interval. (D) Centriculum diameter and SPD-5 area were determine during the development of 1-cell embryos (n=8) expressing SP12::mCherry and SPD-5::GFP (OCF164) at the indicated time points (t=0 is at metaphase). Error bars indicated standard deviation. (E) Representative images of centricula, as detected by SP12::mCherry, in 1-cell embryos at metaphase in a strain also expressing SAS-6::GFP (OCF124) that was treated with the following RNAi conditions: Control RNAi (against *smd-1* (Golden et al., 2009), top row) by feeding, injection of double stranded RNA (dsRNA) again *spd-5* (middle row), and feeding RNAi against *air-1* (bottom row). Control injection of dsRNA against *smd-1* was indistinguishable from feeding RNAi (not shown). Scale bar = 5 μm. (F) Quantification of centriculum diameters of control and *air-1* RNAi as shown in panel (E). Bars represent means and standard deviation. Statistical analysis was done using two-tailed Mann Whitney test. (G) Centriula in 1-cell embryos at metaphase from wild type (OCF176) and *plk-1(or638ts)* (OCF170) worms expressing SP12::mCherry and SPD-5::GFP and grown at the semi-permissive temperature for the *plk-1* mutant. Representative images are shown. Scale bar = 5 μm.

Previous studies showed that the PCM increases in size as cells progress through the cell cycle (Decker et al., 2011). The same is true for the centriculum (Fig. 3D). Moreover, the increase in centriculum size was proportional to the increase in SPD-5 area, as determined at the maximal cross section of the centrosome (the centrosome’s mid-section; Fig. 3D). We thus hypothesized that centriculum size is dependent on the size of the underlying PCM. To test this, we depleted the PCM component, SPD-5, by dsRNA injection into worms expressing the SP12::mCherry ER marker and the SAS-6::GFP centriole marker. This allowed us to determine the location of the centrosome even in the absence of the PCM. Consistent with our hypothesis, SPD-5 depletion led to the disappearance of the centriculum (Fig. 3E, n=14), suggesting that the centriculum requires the PCM for its formation. To further address this possibility, we depleted another PCM component, AIR-1, by feeding RNAi. AIR-1 depletion led to a significant decrease in centriculum size (Fig. 3E, F), although not as dramatically as SPD-5 depletion. It is possible that AIR-1 depletion was not as complete, or because AIR-1 depletion does not affect an already assembled PCM (Cabral et al., 2019). Nevertheless, these results support the notion that the there is a spatial relationship between the PCM and the centriculum. Consistent with this, downregulation of the Polo-like kinase PLK-1 using the conditional *plk-1(or683*^*ts*^) mutant was previously shown to severely affect the membrane in the vicinity of the centrosomes (Rahman et al., 2015). When examining the PCM using SPD-5::GFP in *plk-1(or683*^*ts*^) mutants at the semi-permissive temperature, we found that the centriculum was disorganized but the PCM was still there, albeit smaller (Fig. 3G, n=12). This result further substantiates the conclusion that the centrosome affects centriculum size.

### Increased centriculum size leads to increased PCM size and microtubule nucleating capacity

Given that the centrosome affects centriculum size, it was tempting to explore whether the reverse is also true, namely whether the centriculum affects centrosome size. To date, the only condition that completely abolished the centriculum was by eliminating the SPD-5 PCM component (Fig 3E). However, we did identify a condition, not through inactivation of centrosomal components, that increased the size of the centriculum, allowing us to then ask whether and how this alteration affected centrosome size, and specifically PCM size.

To alter centriculum properties, we tested whether proteins that affect ER organization also affect the centriculum. Specifically, the reticular nature of the centriculum suggested that the dynamin-related GTPase, ATLN-1, the *C. elegans* homolog of atlastin, could play a role in centriculum structure. Atlastin is known for its role in ER-ER fusion (Hu and Rapoport, 2016), and in particular for creating junctions between ER tubules. Tagging endogenously expressed *C. elegans* ATLN-1 with GFP at its C-terminus revealed that it localizes to the endoplasmic reticulum (Supplemental Fig. S3A), similar to the localization of atlastin in other organisms (Chen et al., 2012; Hu et al., 2009; Orso et al., 2009). Consistent with a possible role in centriculum formation, ATLN-1::GFP also localized to the centriculum (Supplemental Fig. S3A). RNAi against *atln-1* led to complete disruption of the ER and embryonic lethality before centricula could be formed (Supplemental Fig. S3B). A milder RNAi treatment (using a mix of *E. coli* expressing double stranded RNA against *altn-1* (20%) and the control *smd-1* (80%)) allowed most embryos to progress through the first mitosis. Under these conditions, centricula in *atln-1* downregulated embryos at metaphase were significantly larger than the control treatment (Fig. 4A and B). However, because the partial RNAi treatment still resulted in embryos that failed to reach mitosis, we also used the auxin-induced degron system (Zhang et al., 2015) that allows for a more precise temporal downregulation of the protein. We created an auxin-inducible *atln-1::degron* construct and treated worms with or without auxin. The requirement of ATLN-1 for cell viability necessitated the use of a short exposure (20-25 minute) to auxin. Nonetheless, this was sufficient to induce a similar change to the one observed by *atln-1* downregulation via RNAi, leading to larger centricula (compare Fig. 4B and Supplemental Fig. S3C). It was previously shown that centrosome size is proportional to embryos size (Decker et al., 2011), raising the possibility that ATLN-1 downregulation affected centriculum size indirectly by affecting cell size. However, the size of control and ATLN-1 downregulated embryos were the same (Supplemental Fig. S3D), excluding this possibility. Thus, downregulation of ATLN-1 lead to an increase in centriculum size.

**Figure 4:**
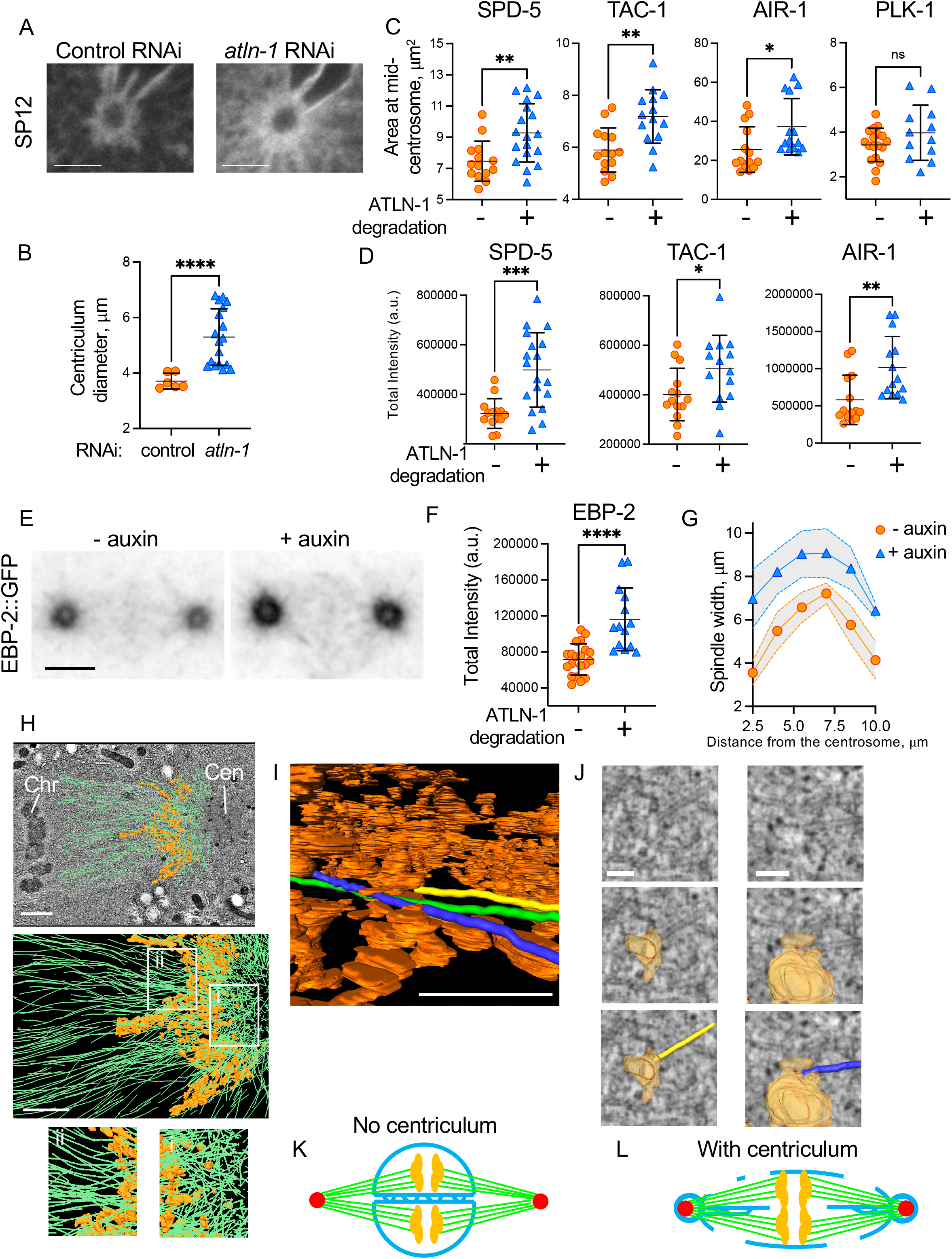
The centriculum affect centrosome function and microtubule organization. (A) Representative examples of 1-cell embryos at metaphase expressing SP12::mCherry (OCF5), following control (left panel) or partial *atln-1* RNAi treatment (right panel). Scale bar= 5 μm. (B) Quantification of average centriculum diameter from the experiment shown in panel A (n=6 and 18 for control (orange) and *atln-1* partial RNAi (blue), respectively). p<0.0001 using the Mann-Whitney test. Error bars here and in panels C, D and F indicate mean and standard deviation. (C) Areas occupied by the indicated fluorescently-tagged centrosomal proteins, as determined at the centrosome mid-plane, in 1-cell embryos at metaphase expressing *atln-1::deg*, without (orange) or with (blue) auxin treatment. The number of centricula analyzed (-/+ auxin) and the p values are as follows: SPD-5 (OCF164): 14/18, p=0.0059; TAC-1 (OCF167): 14/14, p=0.0022; AIR-1 (OCF172): 14/14, p= 0.0106; PLK-1 (OCF166): 18/12, p=0.29. All statistical analyses were done using two-tailed Mann-Whitney test. (D) Intensity of the indicated GFP-tagged centrosomal proteins in the same centricula as in panel C. p values, as determined by two-tailed Mann-Whitney test, are as follows: SPD-5: p=0.0006; TAC-1: p=0.0310; AIR-1: p=0.0035. (E) Localization of EBP-2::mKate2 (OCF162) in a 1-cell embryo at metaphase expressing *atln-1::degron* in the absence (left) or presence (right) of auxin. Scale bar= 5 μm. (F) Quantification of EBP-2 intensity inside the centriculum, as defined by the area inside the SP12::mCherry ring (n= 20 and 14 for - and + auxin, respectively; p<0.001 by two-tailed Mann-Whitney test). (G) Spindle width in the *ebp-2::mKate2*; *atln-1::degron* strain (OCF162), in the absence (orange) or presence (blue) of auxin, as measured at the indicated distanced from one of the centrosomes. Dashed line indicates 95% confidence interval. (H) Top panel: microtubule vectors (green), and a segment of the centriculum (orange) between the centrosome (Cen) and nucleoplasm in a 1-cell embryo in metaphase, superimposed on a single TEM image. Tomography datasets, including microtubule assignments, are from (Redemann et al., 2017). Metaphase chromosomes (Chr) can be seen on the far left of the image. Scale bar= 5 μm. Middle panel: the same microtubule and centriculum reconstructions as in the top panel, but without the TEM image, which obscures structures below the plane of the image. The thickness of this segment, along the z axis, is ∼ 60 nm. Regions i and ii, on the centrosome and chromosome sides of the centriculum, respectively, are enlarged below. Scale bar= 5 μm. (I) Segmentation of three microtubules (marked in yellow, blue and green) shown as they traverse the centriculum (orange). Note that the yellow microtubule terminates at a membrane. See Supplemental Movie S1 for the entire route of these three microtubules through the centriculum. Scale bar = 0.5 μm. (J) TEM images (top row) overlayed with membrane reconstruction (middle row) and membrane + microtubules (bottom row) of the yellow and blue microtubules from panel I and Supplemental Movie S1 at the sites where they hit the membrane. Scale bar= 100 nm. See additional examples in Supplemental Fig. S4C. (K, L) An illustration explaining the possible role of the centriculum in nuclear envelope fenestration during mitosis. Centrosomes are indicated in red, membrane in blue, microtubules in green and chromosomes in orange. See text for further detail.

The increase in centriculum size when ATLN-1 was downregulated allowed us to explore the effect of the centriculum on the centrosome. As noted above, the centrosome is considered a membraneless organelle, and the PCM is assumed to be a LLPS condensate. Therefore, an increase in centriculum size was not expected to influence PCM size. To investigate whether the larger centriculum in ATLN-1 downregulated embryos had any effect on the PCM, the localization of GFP-tagged SPD-5, TAC-1, AIR-1 or PLK-1 was analyzed in 1-cell embryos at metaphase from auxin-treated and untreated worms expressing ATLN-1::degron. ATLN-1 downregulation consistently led to larger centricula for each of these strains (Supplemental Fig. S3C). Unexpectedly, the increase in centriculum size was accompanied by an increase in the area occupied by SPD-5, TAC-1 and AIR-1, but not PLK-1 (Fig. 4C and Supplemental Fig. S3F, G). Moreover, when ATLN-1 was downregulated, the total amount of SPD-5, AIR-1, and to a lesser extent TAC-1, at the centrosome increased (Fig. 4D and Supplemental Fig. S3E). This increase in PCM components may be due to recruitment from cytoplasmic pools as more space becomes available following the increase to centriculum size. The localization and intensity of PLK-1::GFP, which resides in the inner sphere of the PCM (Magescas et al., 2019), remained unchanged when ATLN-1 was downregulated (Supplemental Fig. S3F). Taken together, these results show that increased centriculum area leads to a larger PCM. Thus, despite being a membraneless organelle with a condensate component (Woodruff et al., 2017), the size of the centrosome in *C. elegans* is affected by the membrane that surrounds it.

Finally, we tested whether the change in PCM size due to centriculum enlargement affected the centrosome’s capacity to nucleate microtubules. To do so, we examined the abundance of EBP-2, a plus-end microtubule-binding protein, as a proxy for the number of microtubules along the spindle. Downregulating ATLN-1 caused larger centriculum (Supplemental Fig. S3C and S4A) and led to an increase in the amount of EBP-2 at the centrosome (Fig. 4E, F), suggesting that the increase in PCM size following centriculum enlargement was accompanied by an increased capacity to nucleate microtubules. While the amount of EBP-2 along the spindle was the same with and without ATLN-1 downregulation (Supplemental Fig. S4B), the ATLN-1 downregulated embryos had a wider spindle perpendicular to the centrosome-centrosome axis (Fig. 4G). This is consistent with microtubules radiating from a larger surface, as would be the case for a centrosome that has a larger PCM and surrounding centriculum. This also raised the possibility that the centriculum affects the orientation of microtubule emanating from the centrosome.

### The centriculum may serve as a microtubule “filter”

During mitosis, microtubules originating from centrosomes contact chromosomes for their eventual segregation. Since the *C. elegans* centrosome is surrounded by the centriculum, microtubules must somehow traverse this membrane reticulum to reach the chromosomes. Our FIB-SEM staining protocol was suitable for detecting membranes but less so for microtubules. Therefore, we analyzed the already published vEM tomography data of 1-cell *C. elegans* embryos at metaphase (Redemann et al., 2017). Microtubules in this data set had been traced previously (Redemann et al., 2017), and we added a segmented section of the centriculum between the centrosome and nucleoplasm (Fig. 4H). On the centrosome side, microtubules radiated in all directions (Fig. 4H, middle panel and enlargement i). However, after passing through the centriculum, on the nucleoplasm size, the microtubules were all oriented in the same direction (Fig. 4H, middle panel and enlargement ii). Moreover, the centriculum allowed only a fraction of the microtubules to pass, blocking microtubules that “hit” the membrane (Fig. 4I, J, Supplemental Fig. S4C and Supplemental Movie S1). In the example shown in Fig. 4I and J and Supplemental Movie S1, the microtubules labeled in yellow and blue terminate inside the centriculum, while the microtubule labeled in green pass through the centriculum. In fact, the existence of the centriculum explains the ring-like structures of EBP-2::mKate2 (Fig. 4E and Supplemental Fig. S3H) and tubulin (for example (O’Toole et al, 2012)) around the centrosome. It also explains the preponderance of short microtubules in the vicinity of the centrosome (Baumgart et al, 2019). Thus, the centriculum limits the number of microtubules that are allowed to fully extend. We hypothesize that this results in “microtubule economy”, ensuring that there are sufficient tubulin subunits for a limited number of microtubules to extend all the way to the chromosomes or the cell cortex.

## Discussion

The existence of ER-derived membranes around centrosomes has been known for decades through studies using light microscopy (Bergman et al., 2015; Diaz et al., 2019; Harris, 1975; Karabasheva and Smyth, 2019; Poteryaev et al., 2005; Waterman-Storer et al., 1993), but their configuration and functional importance were unknown. Using vEM methodologies, we show that centrosome-associated membranes in *C. elegans* form a reticulum, leading us to name it the centriculum. The identification of the centriculum underscores the importance of vEM analysis; while the centriculum can be easily detected by conventional EM, in a single imaging plane it appears as an accumulation of vesicles. The reticular nature of the centriculum, on the other hand, provided insight into its possible functions, as discussed below. To our knowledge, this is the first report of a membrane reticulum surrounding a cellular organelle.

Our study sheds new light on the association of the centrosome with the NE, which was shown to be mediated by the LINC complex. The two components of the LINC complex, the Sun domain protein SUN-1 and the KASH domain protein ZYG-12, traverse the inner and outer nuclear membranes, respectively. ZYG-12 was previously shown to localize around the centrosome (Malone et al., 2003), but it was not clear how the LINC complex could do so, given that its components are integral membrane proteins. The presence of the centriculum provides an explanation; indeed, both components of the LINC complex, SUN-1 and ZYG-12, localize to the centriculum (Fig. 1D). It should be noted, however, that unlike the NE, the centriculum does not have obvious membrane structures that are equivalent to an inner or outer nuclear membrane. Thus, the configuration of the LINC complex in the centriculum remains to be established.

In *C. elegans*, centrosomes play a role in nuclear envelope breakdown and the preferential loss of the NPC subunit NPP-3 from the NE adjacent to centrosomes in prometaphase (Hachet et al., 2012; Hachet et al., 2007). We observed that the centriculum is adjacent to the NE in prophase and fused to it in metaphase (Fig. 2 and Supplemental Fig. S1 and S2). It is thus tempting to speculate that the ability of microtubules to enter the nucleoplasm, and loss of NPP-3 from the NE near centrosomes, coincide with centriculum-NE fusion. This step is also reminiscent of the insertion of the fission yeast *Schizosaccharomyces pombe* microtubule organizing center, the spindle pole body, into the NE, which is dependent on the LINC complex (Fernández-Álvarez et al., 2016). Whether the LINC complex is required not only for tethering the centrosome to the NE, as noted above, but also for its insertion once centrosome migration is completed, is currently under investigation.

To fully understand the physiological role of the centriculum we will need to identify conditions that abolish its formation without affecting other pertinent processes, such as centrosome and spindle assembly. Thus far, the only conditions that abolished centriculum formation also disrupted the centrosome (Fig.3E) and/or microtubule assembly (data not shown); the latter was also seen in *Drosophila* (Diaz et al., 2019). Nonetheless, we were able to identify a condition that altered centriculum size, namely the downregulation of atlastin (Fig. 4A-G). How atlastin downregulation affects the structure of the centriculum is currently not known, and attempts to examine the centriculum in *atln-1::degron* strains by FIB-SEM have thus far been unsuccessful due to the fragility of these embryos under high-pressure freezing conditions. Nevertheless, given that centriculum size depends on the PCM, and that PCM size is affected by the centriculum, we speculate that as the centrosome matures there is a balance between outward forces exerted by the growing PCM and its associated microtubules, and resistance applied by the centriculum. When atlastin is downregulated, the centriculum may be less able to resist centrosome-associated forces, leading to an increase in centrosome size.

Regardless of the mechanism by which the PCM expands under conditions of increased centriculum size, the relationship between the PMC and the centriculum introduces a new element when considering centrosome size control. Moreover, this observation suggests that LLPS condensates can be influenced by a confining membrane, which, in itself, should not pose a diffusion barrier to monomeric proteins that make up the condensate. Interactions of a condensate with a membranes are not novel (reviewed in (Zhao and Zhang, 2020)); in many of these cases the membrane serves as a platform for phase separated condensate nucleation. What is unique about our discovery is that the centriculum restricts the size of PCM condensate. In its most simplistic form, the size of a condensate, or the fraction of protein that phase separates to form the condensate, is a function of the properties and concentration of condensate components and the liquid environment in which they reside, such as solute composition and temperature (Shin and Brangwynne, 2017). If that were the case, then increasing centriculum size, which presumably affects neither solute composition nor temperature, should not have affected PCM size. However, not only did the size of the PCR increase, but it also accumulated more PCM proteins, presumably from a soluble pool present in the cytoplasm of the 1-cell embryo. This suggests that the centriculum restricts the size of the PCM, and, hence, its microtubule nucleation capacity. Indeed, the amount of EBP-2 around the centrosome was greater when the centriculum was larger (Fig. 4F).

What might be the function(s) of the centriculum? It was previously suggested that the accumulation of ER around centrosomes in *Drosophila* serves to ensure that daughter cells receive adequate amounts of ER (Karabasheva and Smyth, 2019). We think that this is unlikely to be the case in *C. elegans* given that during mitosis, the ER is distributed throughout the cell and the fraction of the ER around centrosomes is relatively small (Fig. 1A, B). Our study supports several alternative functions for the centriculum: First, the centriculum may limit the number of fully extended microtubules in two ways: it restricts the size of the PCM (Figure 4B), thus limiting its microtubule nucleation capacity, and it blocks a fraction of microtubules from extending past the centriculum (Fig. 4I, J and Supplemental Fig. S4C). These microtubules, presumably, are not in the correct orientation to make it through the entire centriculum. The ability of the centriculum to affect microtubule orientation is also reflected in the wider spindle observe when the centriculum and its underlying PCM are larger due to ATLN-1 downregulation. This could also explain why microtubules appear disordered on the centrosome side of the centriculum, but they become more ordered (namely less overlapping) on the nucleoplasmic side (Fig. 4H). We speculate that had all microtubules nucleated at centrosomes been “allowed” to fully extend, the cell could run out of tubulin subunits before enough astral and kinetochore microtubules reach their targets, the cell cortex and the centromeres, respectively. Thus, the centriculum may serve as a selective barrier that affects both microtubule orientation and number.

Second, the centriculum may be important for tethering the centrosome to the NE during mitosis. As discussed above, the fusion of the centriculum with the NE may facilitate early stages in nuclear envelope breakdown in the vicinity of the centrosomes. Subsequently, NE proteins dissociate from the rest of the NE, but the nuclear membranes remain, becoming highly fenestrated (Rahman et al., 2020). We proposed that the fusion between the centriculum and the nuclear membrane contributes to this subsequent nuclear membrane fenestration as the spindle begins to elongate at metaphase, before sister chromatid separation at anaphase (Velez-Aguilera et al., 2022). Had the centriculum not existed (Fig. 4K), the force generated by the elongating microtubules could not have been transmitted to the nuclear membrane, except by individual microtubules directly attaching to the nuclear membrane, as has been proposed previously (Beaudouin et al., 2002; Salina et al., 2002). The centriculum, on the other hand, provides a mechanism for transmitting the force generated by microtubule elongation to the nuclear membrane (Fig. 4L): as the spindle elongate and the centrosomes move apart, the centrosomes pull the centricula, which, in turn, pull on the rest of the nuclear membrane. Assuming that there is no net increase in membrane at this stage, this pulling action will cause the membrane to fenestrate, as is observed during NE breakdown.

In summary, in this study we identify a centrosome-associated membrane reticulum, the centriculum, and show that there is a mutual dependency between the centriculum and the centrosome: the centriculum depends on the presence of the centrosome, and the size of the PCM is affected by the size of the centriculum. Our data suggested that the centriculum also affects centrosome function by orienting microtubules and limiting how many can fully elongate. Given the conservation in centrosome components and ER organization, centricula are likely to exist in other organisms.

## Acknowledgements

We thank Alexander Dammermann (University of Vienna, Austria), Lionel Pintard (institut Jacques Monod, France), Jessica Feldman (Stanford University, USA), Jon Audyha (University of Wisconsin-Madison), Geraldine Seydoux (Johns Hopkins University School of Medicine) and the Caenorhabditis Genetics Center (CGC, University of Minnesota, USA) for strains and reagents, and Tetsunari Fukushige for help with generating dsRNA. We also thank Will Prinz and Kevin O’Connell for valuable discussion and comments on the manuscript. R.M., M.M.R., S.D., M.O. and O.C.-F. were funded by an Intramural NIDDK grant number DK069012. G.F. and T. M.-R. were funded by the German Research Foundation (DFG grant MU 1423/8-1 and 8-2). M.A.Q.M. and D.Q.M. were funded by the NCI (F30CA257383) and NIGMS (R01GM121597).

## Author contribution

R.M., M.M.R, S.D, M.O, D.Q.M. and O.C-F designed the experiments. R.M., M.M.R, S.D., M.O and M.A.Q.M. carried out the experiments. All authors were involved in data analysis. M.M.R. and S.D. carried out the segmentation of the FIB-SEM data. M.M.R., S.D., G.F. and T. M-R. were involved in analyzing the EM tomography data. The first draft of the manuscript was written by R.M., M.M.R, S.D. and O.C-F. All authors reviewed and edited the manuscript drafts. The authors declare no competing interests.

## Materials and Methods

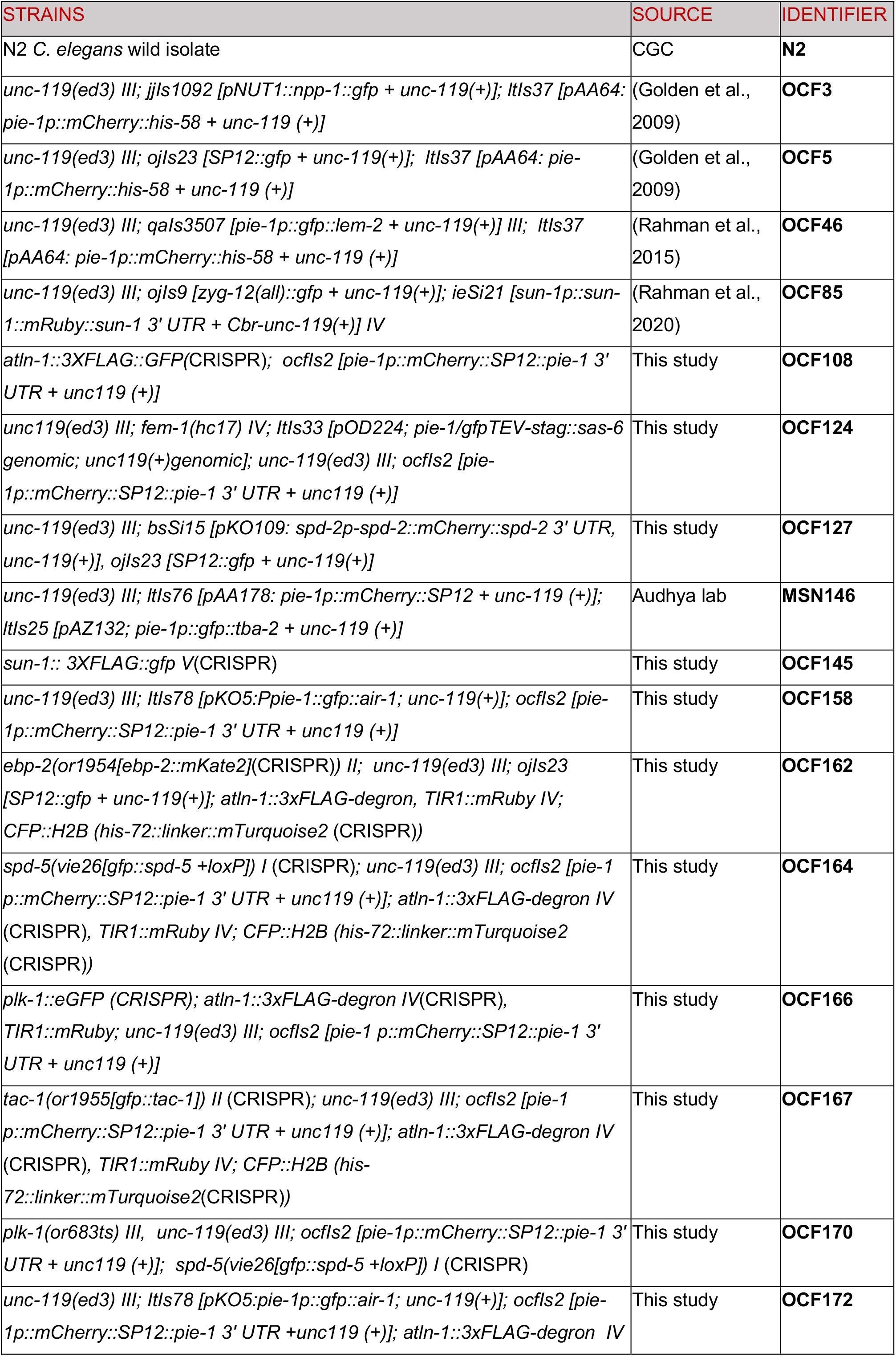
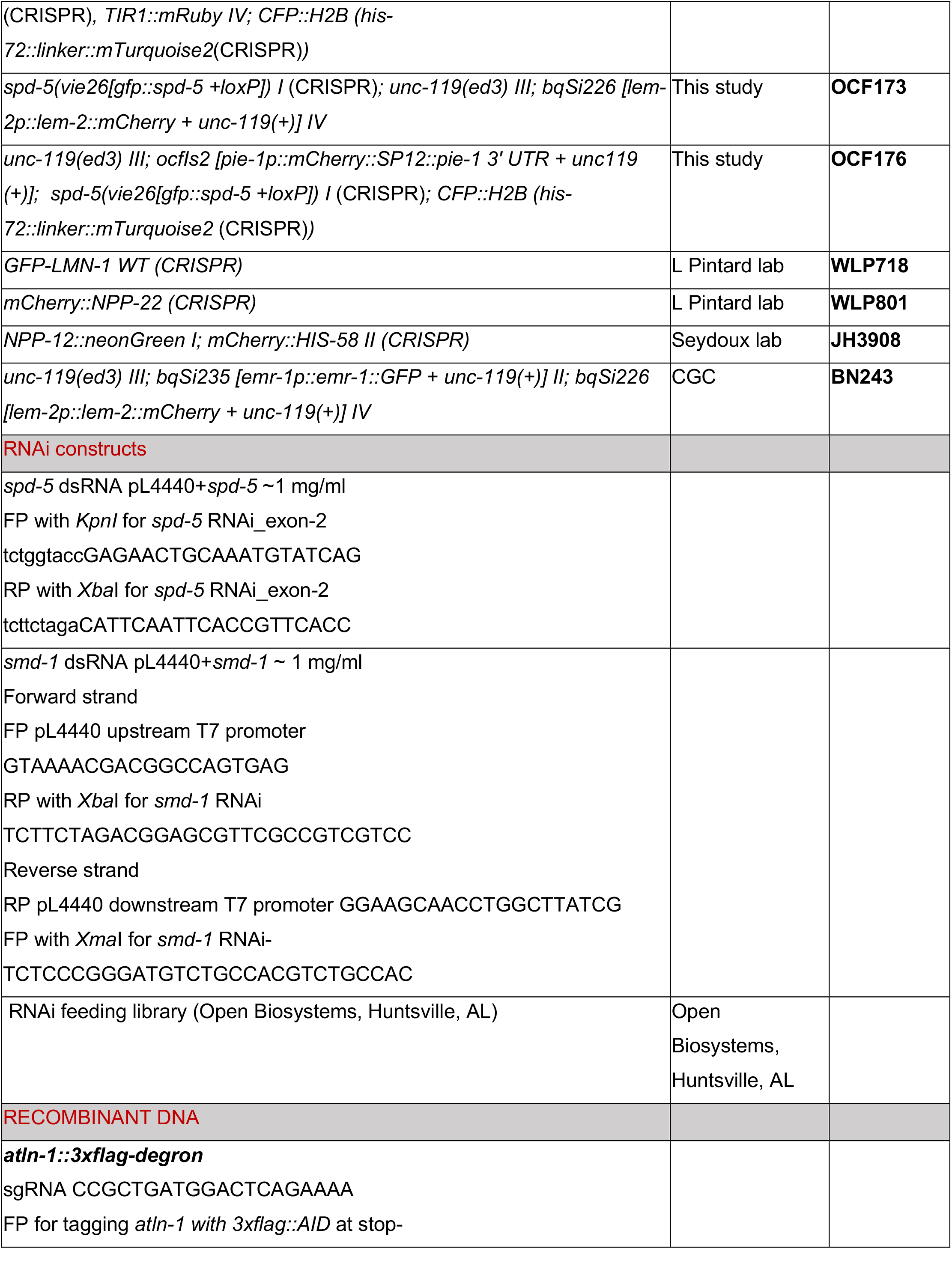
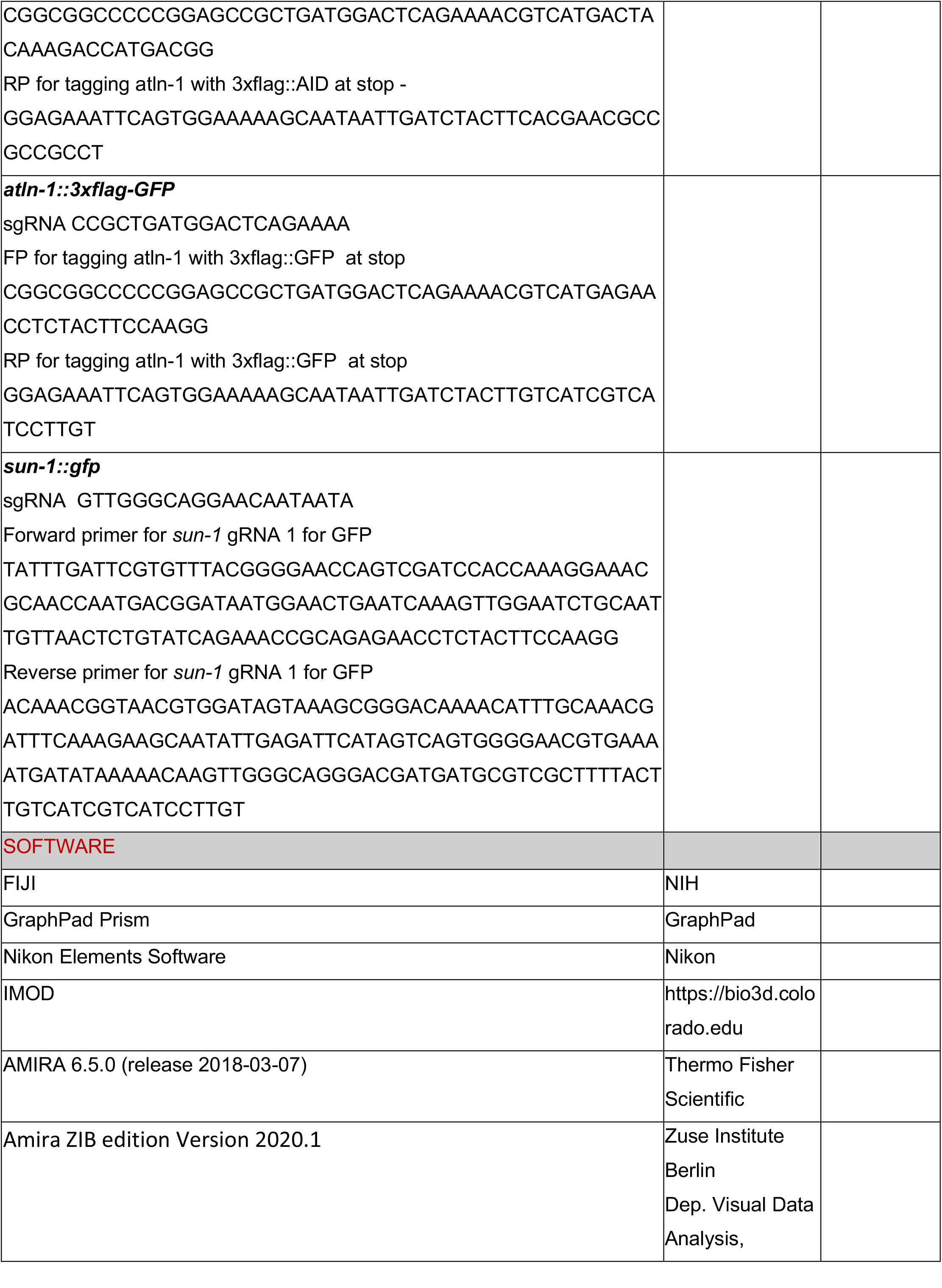
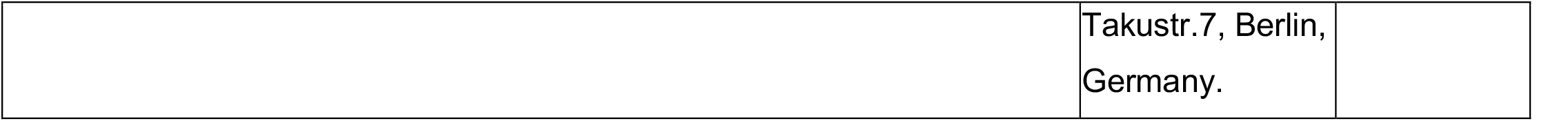

## EXPERIMENTAL MODEL DETAILS

*C. elegans* strains expressing mCherry::SPD-2 (Peel et al., 2017), SAS-6::GFP (Dammermann et al., 2008), PLK-1::sGFP (Martino et al., 2017), GFP:AIR-1 (Portier et al., 2007), GFP ::TAC-1 (Chuang et al., 2020), EBP-2::mKate2 (Sugioka et al., 2018), GFP::SPD-5 (Cabral et al., 2019), CFP::H2B (Sloan and Bembenek, 2021), SP12::GFP, SP12::mCherry (Joseph-Strauss et al., 2012), EMR-1::GFP (Morales-Martínez et al., 2015), NPP-1::GFP (Golden et al., 2009), LMN-1::YFP (Galy et al., 2008), LEM-2::GFP (Morales-Martínez et al., 2015), LEM-2::mCherry (Morales-Martínez et al., 2015), ZYG-12::GFP (Malone et al., 2003), and the temperature sensitive mutant *plk-1 (or683ts)* (O’Rourke et al., 2011) have been described previously. Strains expressing endogenously tagged ATLN-1::GFP, 3XFlag::degron tagged ATLN-1 and SUN-1::GFP were generated by CRISPR/Cas9-induced homologous recombination (Dickinson et al., 2013).

## METHOD DETAILS

### RNA-Mediated Interference

#### Feeding RNAi

For *air-1* feeding RNAi, a 5 ml Luria Broth (LB) with 50 mg/ml ampicillin were inoculated using 1:100 inoculum from a 2 ml overnight saturated culture (at 37 °C) of *E. coli* carrying a plasmid with either *smd-1* (control RNAi) or *air-1* expressed from both ends (RNAi clones are from the RNAi feeding library (Open Biosystems, Huntsville, AL). Once the culture grew to OD_600_ of around 0.5 (∼ 4 h at 37 °C), 0.5 M IPTG was used to induce the bidirectional transcription of *air-1* (1 mM final concentration) for another 4 h. The culture was centrifuged at 5000 g for 5 min at room temperature, and the pellet was resuspended in 1 ml of fresh LB + ampicillin (50 mg/mL) media. 200 μL of this culture was spread on each RNAi plate (MYOB with 4 mM IPTG and ampicillin 50 mg/mL). For feeding RNAi treatment, 15-20 L4-stage larvae were transferred to RNAi plates, and after 48 h, the RNAi treated worms were dissected on a glass slide (Cat #EF15978A, Daigger Scientific), mounted on a 2% agar pad and the early embryos were imaged as described below. For embryonic viability scoring, a few worms were allowed to lay embryos for 3-4 h on a fresh RNAi plate, and embryonic viability was scored after 24 h.

#### Injection of dsRNA

For dsRNA against *spd-5*, approximately 1 kb of the coding region was cloned in plasmid pL4440 (Addgene). The plasmid containing the *spd-5* coding DNA was linearized at one end of the coding region using KpnI restriction enzyme and subsequently cleaved by XbaI. For dsRNA against *smd-1*, around 1 kb was PCR amplified from pL4440+*smd-1* plasmid from the RNAi feeding library (Open Biosystems, Huntsville, AL). For forward strand, a forward primer upstream of the first T7 promoter site and a reverse primer at the end specific to the *smd-1* were used. Similarly, for the reverse strand, a reverse primer downstream of the T7 promoter site at the other end and a forward primer at the beginning specific to the *smd-1* were used. Digested or PCR amplified fragments were purified using the Qiagen MinElute Reaction Cleanup Kit (Cat# 28206, Qiagen). To prepare dsRNA for injection, *in vitro* RNA synthesis was carried out using “MEGAscript™ T7 Transcription Kit (Cat# AM1333, Invitrogen) followed by purification using Phenol:CHCl_3_:Isoamyl Alcohol (25:24:1, v/v) (Cat #15593031, Invitrogen) and precipitated using 100% ethanol (Cat #64-17-5, The Warner-Graham company). The RNA pellet was dissolved in ∼100 to 150 μL of TE buffer (10 mM, pH 8) (Cat #351-011-131, Quality biological). To prepare dsRNA, about 100 μl (1 μg/μl) of the ssRNA was mixed and kept at 85 °C for 3 min in an aluminum heat block incubator followed by slow cooling to RT for annealing. Injection of dsRNA was done according to Ohta laboratory protocol (Ohta et al., 2021). L4s (15-20 worms) were injected with ∼1 μg/ μL dsRNA. These worms were maintained at 16 °C for 48 h prior to live imaging of early embryos by confocal microscopy.

#### Downregulation of ATLN-1 using auxin-induced degradation

For auxin mediated degradation, the protein of interest is tagged with an auxin-inducible degron tag, and cells also expressed TIR1, an exogenous F-box protein (Zhang et al., 2015). One of two types of controls was used: either worms with or without TIR1, both treated with auxin, or ATLN-1::degron TIR1 worms with and without auxin treatment. Worms were transferred to bacteria seeded auxin plates (MYOB plates with 4mM auxin analog, indole-3 acetic acid (IAA), Alfa Aesar, #A10556) for 20-25 minutes, and embryos were imaged immediately thereafter. Auxin plates (4 mM) were prepared from a 400 mM auxin stock (in 100% ethanol and stored at 4°C for up to a month). Auxin was added to the MYOB media before pouring plates at ∼50°C. Next day, auxin plates were seeded with a thick grown OP50 culture and left to dry in a dark place. Based on (Zhang et al., 2015).

### Confocal microscopy and image processing

Unless indicated otherwise, images were taken using a Nikon confocal Ti2 with Yokagawa CSU-X1 spinning disk and a photometrix Prime 95B camera using a Nikon water/oil 60X 1.2-NA Apo Plan objective. Images were captured using Nikon Elements software version 5.20.00. Images in Fig 1C were taken using a a custom-assembled spinning disk confocal microscope consisting of a Zeiss Axio Imager A2 frame, a Borealis modified Yokogawa CSU10 spinning disk, an ASI 150-micron piezo stage controlled by a MS2000, an ASI filter wheel and a Hamamatsu ImagEM X2 EM-CCD camera using a Plan Apochromat 100x/1.4 NA DIC objective (Carl Zeiss). Images in Figure 4A were taken using a Nikon confocal Ti2 with Yokagawa CSU-X1 spinning disk with a Prime BSI camera, using a 60X Plan Apo VC objective, NA 1.2. Images in Fig. S3B were taken using Nikon Eclipse TE2000U spinning-disk confocal microscope with Metamorph software version 7.8.6.0 (Molecular Devices). The microscope was equipped with a 60X 1.4-NA Apo objective, an LMM5 laser merge module with four diode lasers (excitation at 405, 491, 561, and 655 nm) from Applied Research, a Yokogawa CSU10 spinning disk, and a Hamamatsu C9100-13 EM-CCD camera. For imaging, embryos were mounted on 2% agarose (Cat #214010, Invitrogen) pads prepared in standard M9 buffer (Cat# 11006-517, IPM scientific). Images were taken at z = 1 μm intervals unless otherwise mentioned. Images were processed with Fiji (ImageJ release 2.1.0; http://imagej.nih.gov/ij) or Adobe Photoshop CC (release 23.1).

## QUANTIFICATION AND STATISTICAL ANALYSIS

### Image analysis

All images were analyzed using Fiji (ImageJ release 2.1.0; http://imagej.nih.gov/ij). For centriculum diameter, two perpendicular lines (width 7 μm) were drawn across the central plane of the centriculum, and intensity profile of the plotted line was obtained. The two peaks of maximal intensity represented the two middles of the centriculum ring width as the line traversed the centriculum. A rectangle tool was used to calculate the distance between the two peaks, which gave us two diameters (d1 and d2) of the centriculum. The diameters were then used to determine the average diameter (d1+d2)/2 of the centriculum.

For embryo size, a freehand selection tool in FIJI was used to trace the edges of the central plane of the one-cell embryos at metaphase stage. The software provided the area of the traced region.

To determine PCM size, we measured the area of the PCM in a maximal cross section (the central plane of the centrosome) using the trace and thresholding method in Fiji. The mode pixel intensity was determined and subtracted from the intensities at the z-slice of interest. A threshold was applied, which remained constant for both control and test, followed by drawing a circle around the thresholded area, which gave us the area and raw intensity of the thresholded region.

For PLK-1 intensity, a line was drawn across the PLK-1::sGFP occupied area at the central plane of the centrosome, and an intensity profile was plotted. For AIR-1, TBA-2, and SP12 localization graphs, a 9 μm line (line width = 1 μm) was drawn through the centrosome perpendicular to the spindle. The intensity profile was plotted for the ER channel (SP12::mCherry) and the protein of choice channel (AIR-1::GFP and TBA-2::GFP respectively). The average values across the line for both channels were plotted.

For EBP-2::mKate2 intensity inside the spindle, a 10 × 1 μm wide rectangular box drawn perpendicular to the long spindle axis was used to measure the integrated density of fluorescence at 2 μm and 4 μm away from the center of one of the centrosomes.

Strain OCF162 expressing EBP-2::mKate2 was used to determine the spindle width. The height of the spindle was measured by placing vertical lines (width 7) at the following intervals from the center of the centrosome: 2.5 μm, 4.0 μm, 5.5 μm, 7.0 μm, 8.5 μm, and 10 μm. For each line, an intensity profile of EBP-2::mKate2 was generated. From the intensity profile, the top and bottom of the spindle was identified, and the height of the spindle was measured between these two sites.

For imaging adult somatic cells, we used a custom-built spinning disk confocal microscope with a 100x oil objective (Adikes et al., 2020). First, worms at the L3 larval stage were immobilized using 5 mM levamisole (Cat# L9756, Sigma-Aldrich) in M9 buffer. Subsequently, the immobilized worms were mounted on a 5% Noble agar (Cat #A5431, Sigma-Aldrich) pad split into two asymmetric halves with worms placed on the larger half. A coverslip (Cat #12-541B, ThermoFisher Scientific) was gently placed on the worms. We used VALAP (http://cshprotocols.cshlp.org/content/2015/2/pdb.rec082917) to secure the coverslip onto the microscope slide, leaving two diagonal openings. We flooded the chamber with M9 buffer to prevent desiccation. Finally, we set the step size at 1 μm and imaged every 2.5 minutes for 3 hours using 488 and 561 nm lasers.

### Statistical analysis

Mann-Whitney, one-way ANOVA or Student’s t-tests were conducted using GraphPad Prism [version 9.1.2 (255)], as indicated in the relevant figure legends. The number of samples analyzed is indicated in the text or figure legend. P-values are indicated in the respective figure legends.

### Temperature Shift Experiments

For temperature shift experiments, embryos from *plk-1*^*ts*^ animals expressing SPD-5::GFP; SP12::mCherry were maintained at 16°C. *plk-1*^*ts*^ homozygous adult hermaphrodites were shifted to 23°C and the early embryos were imaged after 30-60 min incubation.

### Focused Ion Beam-Scanning Electron Microscope (FIB-SEM)

#### Embryo collection and sample processing

We dissected wild-type N2 hermaphrodites (maintained at 20°C) in 20% Bovine Serum Albumin (BSA) and collected single 1-cell embryos in a cellulose capillary tube (Pelletier et al., 2006) (Cat# 16706869, Leica Microsystems, Vienna, Austria) under a stereomicroscope (SMZ645, 50X total magnification, Nikon). We followed the zygotes under the same stereoscope until they reached the desired stage in mitosis, i.e., prophase, prometaphase, metaphase. Then we froze the embryo immediately using gold-coated planchettes (Cat #16770152, Cu-Au 3.0 × 0.5 mm, Leica Microsystems, Vienna, Austria) using a high-pressure freezer (model EM ICE; Leica Microsystems, Vienna, Austria). We then applied quick-freeze substitution (QFS) (McDonald and Webb, 2011) with a cocktail of 0.2 g Osmium tetroxide (OsO_4_) powder (Cat# 19100, Electron Microscopy Sciences in 9 ml of acetone (Cat #10000, Electron Microscopy Sciences), 0.5 ml of double deionized water, and 0.5 ml of 2% uranyl acetate (UA) solution (Cat #22400, Electron Microscopy Sciences). Then we washed the samples for 10 minutes (x3) with 100 % acetone at room temperature and infiltration in Polybed 812 resin (Cat #08791; Polysciences: 14.6 g Polybed, 8.4 g DDSA, 7.0 g NMA, and 0.42 ml DMP30) followed by resin: acetone ratios of 1:2, 1:1, and 2:1 each for an hour. Finally, we transferred the samples into 100% resin for ∼16 hours prior to curing at 55°C oven for 40 to 60 hrs. The cured resin was trimmed using a jeweler’s saw and razor blades and sectioned in an ultramicrotome until the embryos were tangentially exposed for FIB-SEM imaging. For a step-wise protocol, please refer to (Rahman et al., 2021).

#### FIB-SEM data acquisition and postprocessing

Prior to imaging, the embryos were sputter coated with a 10 nm carbon coat to increase overall conductivity, and a thicker ∼300 nm carbon pad was deposited with the FIB to cover the buried embryo. FIB-SEM acquisition was done using a Crossbeam 540 (Carl Zeiss). The front edge of a sample was milled with FIB beam (at 65 or 30 nA) followed by a reduced FIB current (3 nA). Samples were milled until the embryos were detected at the cliff face. At this point, a 1.5-μm-thick patterned platinum and carbon pad was laid over the carbon pad (Narayan et al., 2014). Milling with a FIB current of 1.5 nA was resumed across the entire embryo, with intermittent SEM imaging, until a recognizable biological feature such as the hazy pericentriolar material was revealed. Next, imaging was initiated using ATLAS 5 (Fibics) at high-resolution for a region of interest in the embryo. The SEM was operated at 1.5 kV accelerating voltage and 1.0 nA current. FIB parameters were 30 kV with a 1.5 nA or 700 pA current. High-resolution, high-m/z contrast signal was recorded using an in-column energy selective back-scatter detector with a grid voltage of 1000 V. Images at 3 nm pixel size, dwell time of 3 μs, and 9 nm FIB step size or “z thickness” were obtained. Several thousand high-resolution images were captured in 40-60 h. After dataset acquisition, the image stack was cropped and processed using IMOD-based scripts (Kremer et al., 1996) to produce aligned, inverted, and binned mrc-image volumes with 9 nm isotropic voxels.

#### Segmentation of FIB-SEM datasets to generate 3D models

We used Amira 6.5.0 (release 2018-03-07; Thermo Fisher Scientific) software with XIMagePAQ, XMesh, and XSkeleton extension packages for a threshold-based semi-automated segmentation. Segmentation using Amira was done following a general scheme unless specified elsewhere. We made a threshold-based selection of membranes first, followed by three rounds of slice-by-slice visual inspection (through XY, YZ, and XZ planes separately). Upon careful visual inspection, we manually added all undetected areas (about 1-2% of the total area) to the segmented volume. We removed any unrelated structures that were automatically selected due to a similar threshold level, (e.g., ER membranes beyond centriculum; nuclear envelope, lipid droplets, mitochondria, and other organelles in the proximity of centriculum). In our current study, we included FIB-SEM data from four wild-type embryos: two prophase (P1 and P2), and two metaphase (M1 and M2) embryos.

#### Segmentation of FIB-SEM datasets to generate 3D models

The software Amira ZIB (Zuse Institute Berlin) version 2020.1 was used for manual segmentation of membranes around centrosome. Segmented membranes were visualized following the Amira default scheme unless mentioned otherwise. Segmented membrane sections were inspected twice slice-by-slice through visual inspection of the XY, YZ, and XZ planes, separately. Any undetected area (about 1-2% of the total area) left during segmentation was manually added to the final volume. Any unrelated area, for example, round and isolated vesicles less than 300 nm in diameter (Hammerling et al., 2020), were removed manually.

### Electron tomography of serial sections

#### Embryo collection and sample processing

We used samples that were previously obtained for a 3D reconstruction of the first mitotic spindle in *C. elegans* (Redemann et al., 2017). Briefly, *C. elegans* N2 (wild type) gravid adults were dissected in M9 buffer, and zygotes in early mitosis were collected in cellulose capillary tubes (Cat# 16706869, Leica Microsystems) (Pelletier et al., 2006).

The embryos were observed under a stereoscope until metaphase was reached and then immediately frozen using an EMPACT2 high-pressure freezer equipped with a rapid transfer system (RTS, Leica Microsystems). Freeze substitution of the cryo-immobilized embryos was done over three days in anhydrous acetone containing 1% OsO4 and 0.1% UA using freeze-substitution equipment (EM AFS, Leica Microsystems, (Pelletier et al., 2006)). Epon/Araldite infiltration was followed by thin-layer-embedding and polymerization for three days at 60°C. After remounting the specimens on dummy blocks, serial semi-thick sections (300 nm) were cut using an ultramicrotome (Ultracut UCT, Leica Microsystems). Sections were collected on a Formvar-coated copper slot grids (EMS) and post-stained with 2% UA (in 70% methanol) followed by brief exposure to Reynold’s lead citrate (Müller-Reichert et al., 2007).

#### Data acquisition by electron tomography and postprocessing data analysis

Prior to imaging, 15 nm colloidal gold particles (Cat# 777137, Sigma-Aldrich) were added to both sides of the 300 nm semi-thick sections to serve as fiducial markers for the calculation of electron tomograms. Next, a series of tilted images was taken by using a TECNAI F30 transmission electron microscope (Thermo Fisher Scientific) operated at 300 kV. TEM images were taken every 1° over a ±60° range using a Gatan US1000 CCD camera (2k X 2k). The pixel size of the images was 2.3 nm. Tilted views/images were aligned using the fiducial gold markers and later back-projected and combined to a super montage using IMOD software package (Kremer et al., 1996). To cover the pole-to-pole region of each mitotic spindle, on average, 24 consecutive serial sections per spindle were imaged and processed accordingly (Redemann et al., 2017).

The Amira 6.5.0 (release 2018-03-07; Thermo Fisher Scientific) software with XIMagePAQ, XMesh, and XSkeleton extension packages was used for microtubule segmentation. Microtubules were segmented in individual sections prior to serial stitching based on a previously published template matching and stitching algorithms (Lindow et al., 2021; Weber et al., 2011). Segmented vector datapoints (in tomogram sections #6 and #7) for MTs from the same tomogram dataset (Redemann et al., 2017).

