## Supplemental Figures for "The *Caenorhabditis elegans* centrosome is surrounded by a membrane reticulum, the centriculum, that affects centrosome size and function"

#### Supplemental Materials

#### Supplemental Figure Legends

**Supplemental Figure S1: Centricula in prophase embryos**

(A, B) Two centricula were segmented (orange) from 1-cell embryos at prophase as described in the main text and in the legend to Figure 2. (i, ii) a single SEM plane with (i) or without (ii) a superimposed 200 nm segmented centriculum section from planes above the SEM image shown. Arrows point to the nuclear envelope. (iii) An 800 nm slice of the centriculum. (iv) A view through the entire centriculum wall between the centrosome (forefront) and the pronuclei (background). Scale bar = 1 µm except panel iv, where scale bar = 100 nm.

**Supplemental Figure S2: Centricula in metaphase embryos**

(A, B) Two centricula were segmented (orange) from 1-cell embryos at metaphase. The panels are as described in the legend for Supplemental Figure S1. Scale bar = 1 µm except panel iv, where scale bar = 100 nm.

**Supplemental Figure S3: The centriculum affects centrosome structure and function.** (A) A 1-cell embryo (strain OCF108) expressing SP12::mCherry (left panel) and endogenously tagged ATLN-1::GFP (center panel), which co-localizes with SP12 (merged image, right). Scale bar= 10 µm. (B) The effect of ATLN-1 downregulation by feeding RNAi. Worms expressing SP12::GFP and histone H2B::mCherry (OCF5) were exposed to RNAi against *atln-1* for 48 hours and embryos were imaged. Scale bar = 10 µm. (C) Quantification of the effect of auxin treatment on centriculum size in 1-cell embryos expressing *atln-1::degron* and the indicated fluorescently-tagged centrosomal proteins, without (orange) or with (blue) auxin treatment. Measurements were done on 1-cell embryos at metaphase. The number of centricula analyzed (-/+ auxin) were as follows: SPD-5 (OCF164): 14/18; TAC-1 (OCF167): 14/14; AIR-1 (OCF172): 14/14; PLK-1 (OCF166): 18/12; EBP-2 (OCF162): 20/14. Note that these data are from the same experiments described in Figures 4C-F. Error bars here and in all subsequent graphs in this figure indicate mean and standard deviation. All p values <0.0001, as determined by one-way ANOVA. (D) The size of 1-cell embryos taken from worms expressing SPD-5::GFP, SP12::mCherry, CFP::H2B and *atln-1::degron* (strain OCF164), with and without auxin treatment, was measured as described in the Material and Methods section. n= 7 and 9 embryos for – and + auxin treatment, respectively. p=0.6065, as determined by Mann Whitney test. (E) Localization of the indicated centrosome proteins fused to GFP in strains also expressing *atln-1::degron*, as described in the legend of Figure 4C, in the presence or absence of auxin. The quantification of the area occupied by these proteins is shown in Figure 4C. (F) Fluorescence intensity of PLK-1::GFP along a line that traverses the center of the centrosome in strain OCF166, also expressing *atln-1::degron*. Dashed lines indicate 95% confidence interval.

**Supplemental Figure S4: The centriculum serves as a microtubule filter.**
(A) the Localization of EBP-2::mKate2 and SP12::GFP (strain OCF162) in 1-cell metaphase embryos also expressing ATLN-1::degron, in the presence or absence of auxin. Scale bar= 5 µm. (B) EBP-2::mKate2 intensity in metaphase 1-cell embryos also expressing *atln-1::degron* (OCF162) along the spindle at 2 and 4 µm from the centrosome, in the absence (orange) or presence (blue) of auxin. The number of centrosomes analyzed was 22 for the – auxin and 14 for + auxin. Comparison between the two conditions at the indicated distances was done using one-way ANOVA. p values in both cases were 0.9469. (C) Three examples of microtubules (in yellow) terminating at membranes (orange). Top row: a single TEM image. Middle row: 3D reconstruction of a membrane segment where a microtubule terminates. Bottom row: 3D reconstructions of both the microtubule and the membrane. Note that the reconstructions use data from tomography images adjacent to the ones shown in the top row. Scale bar= 100 nm.

**Supplemental Movie S1**

3D reconstruction of the microtubules and membrane shown in Figure 4I. The movie, done using Amira software, tracks the microtubules as the pass through the centriculum. The yellow and blue microtubules end at the membrane, while the green microtubule goes all the way through.

Supplemental Figure S1

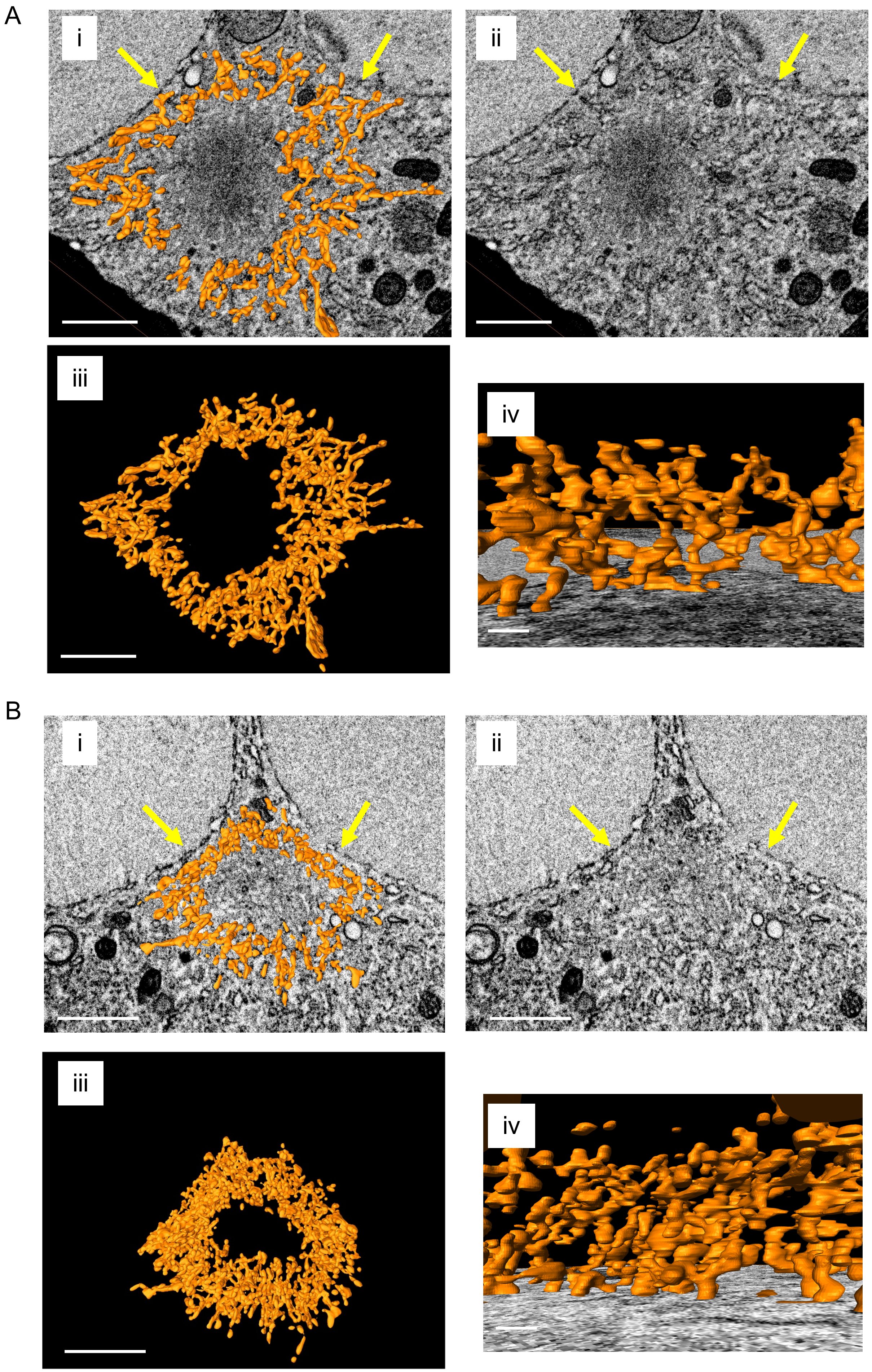

### Supplemental Figure S2

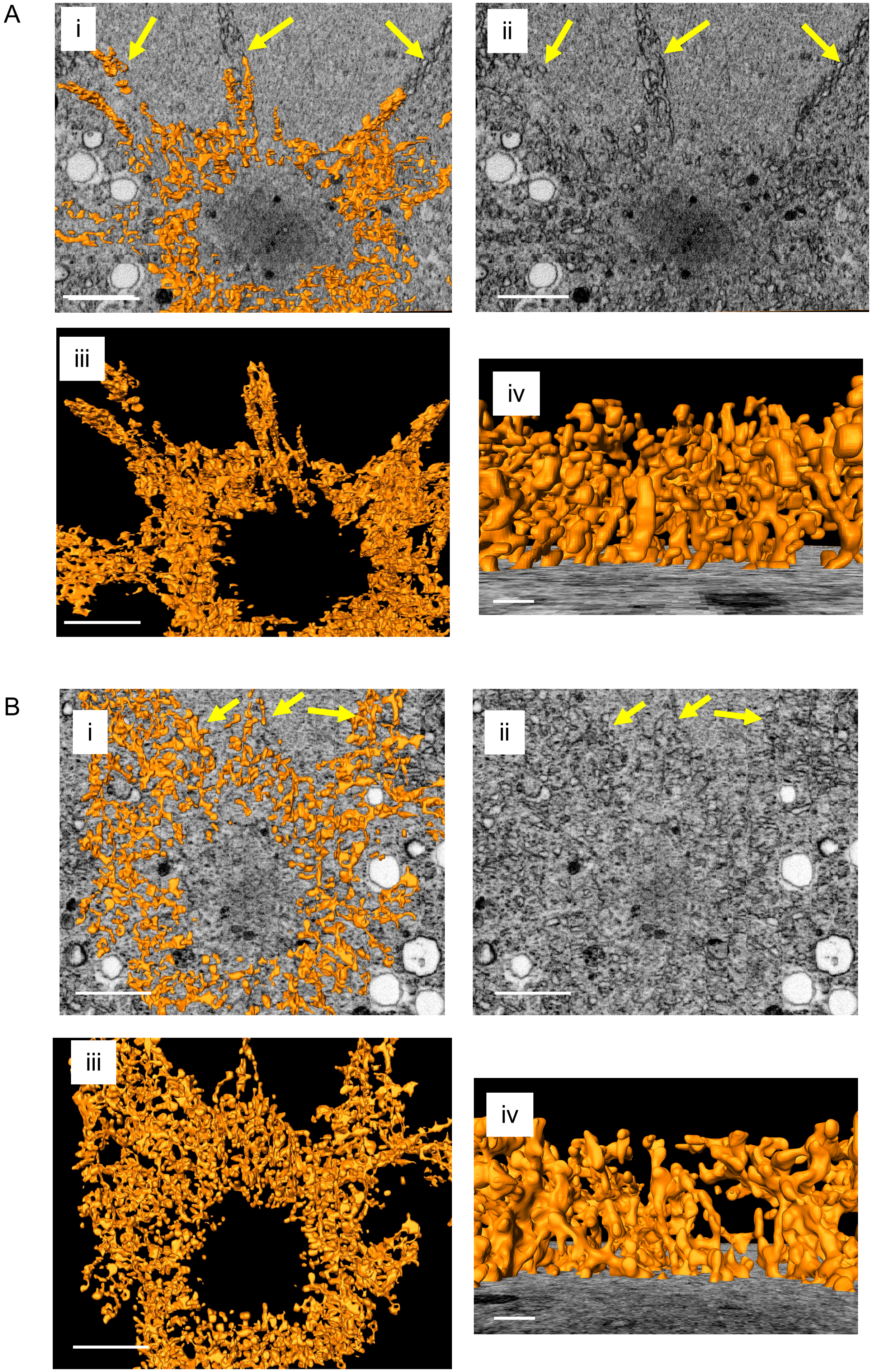

### Supplemental Figure S3

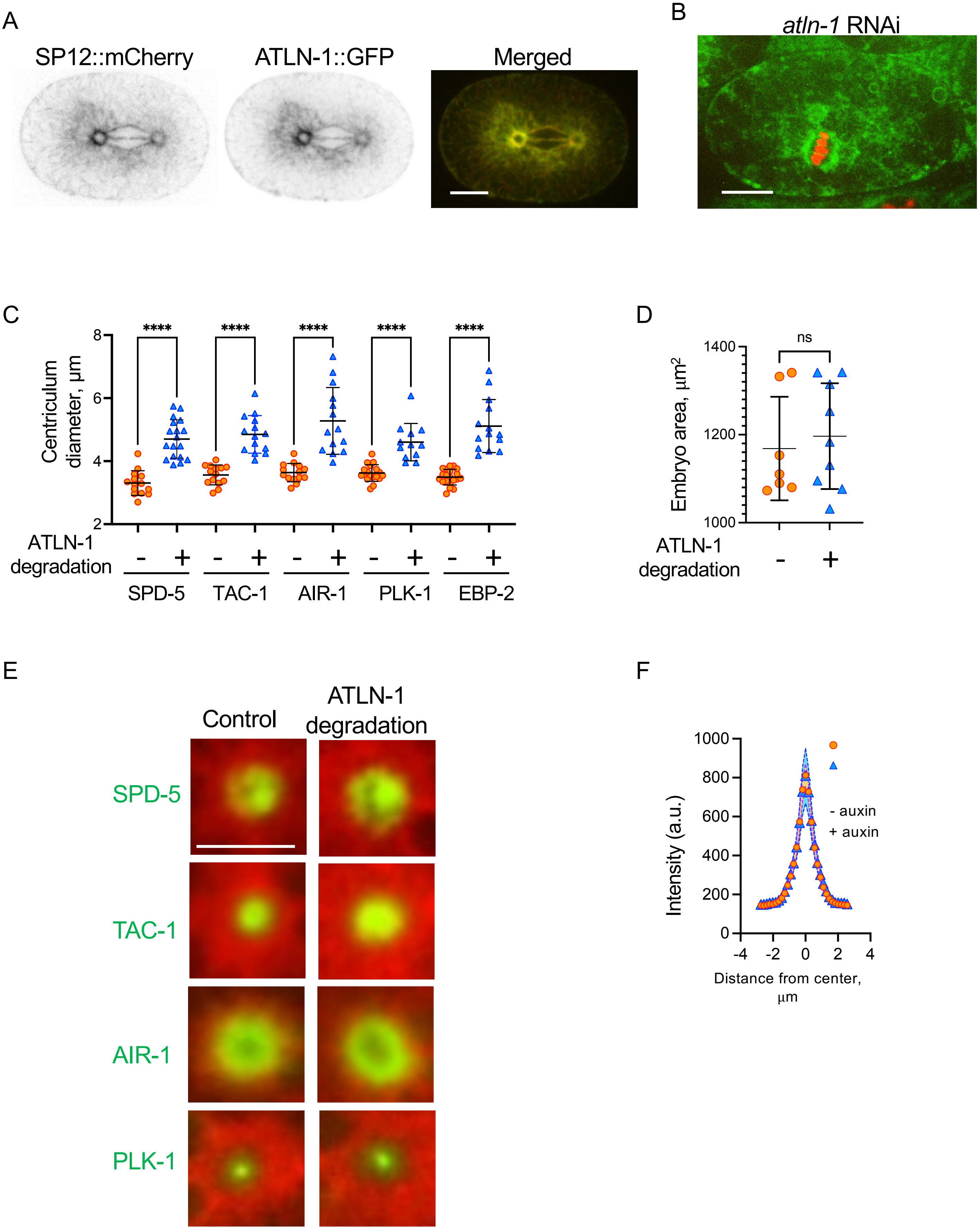

### Supplemental Figure S4

A

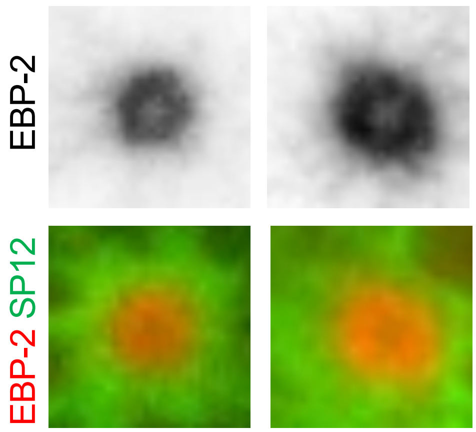


B

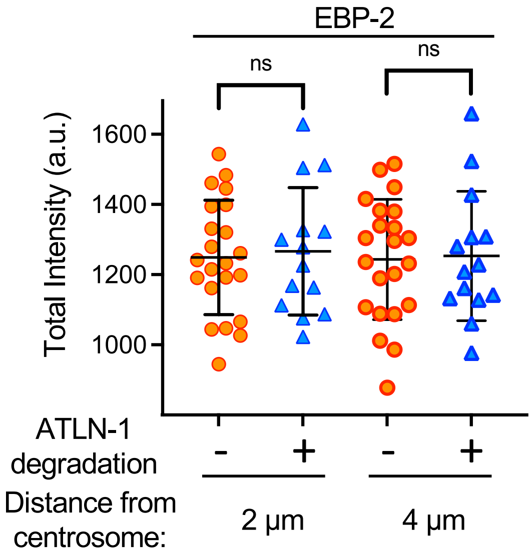


C

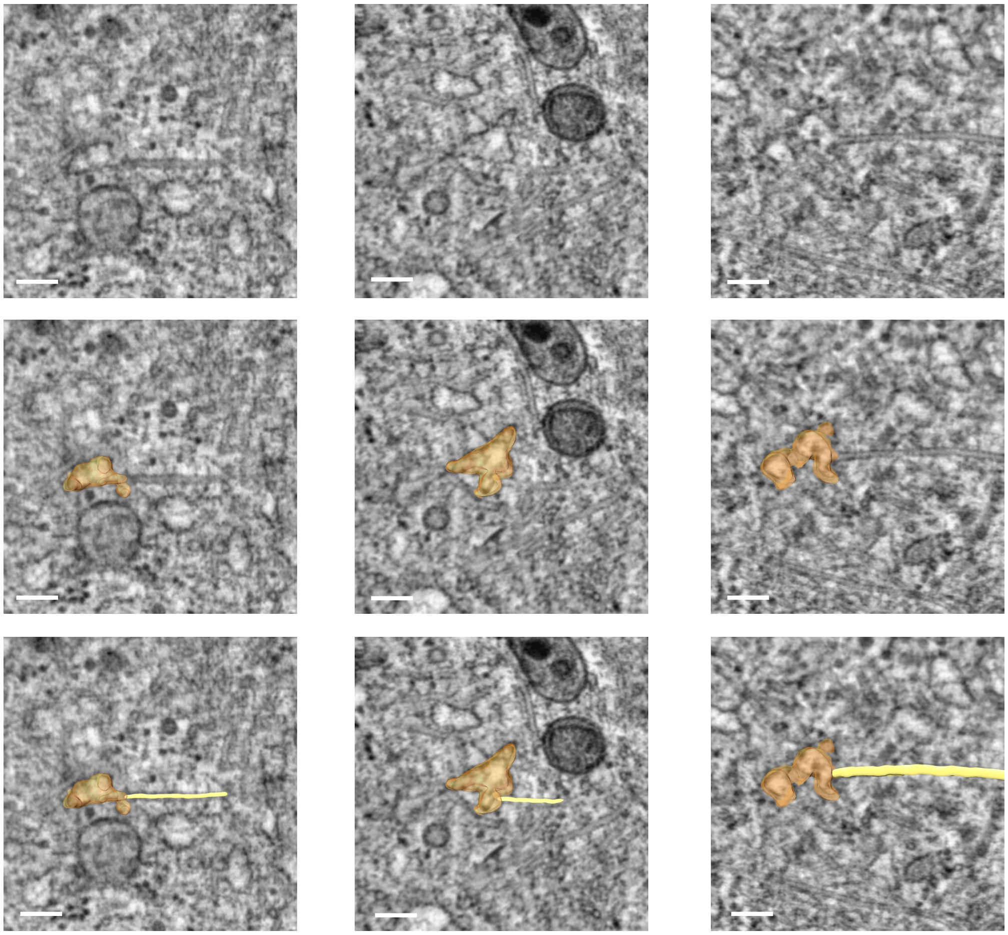
